# Speed judgments during smooth pursuit reflect low gain efference copy

**DOI:** 10.64898/2026.10.29.762385

**Authors:** Shenyue Zhao, Anamika Sudheer, Maureen Ann Hagan, Nicholas Seow Chiang Price

## Abstract

During smooth pursuit eye movements, the visual system must compensate for self-generated retinal motion to achieve stable perception of motion in the environment. This compensation primarily depends on an efferent copy of motor commands. Systematically under- or overweighting the strength of efference copy should lead to correlations between eye movement velocity and perceived object velocity. However, previous examinations of these trial-by-trial perceptual correlations have produced mixed results. It remains unclear how effectively the brain compensates for eye movement induced retinal motion during pursuit, and what this reveals about the way efference copy influences perception. To address this, we developed a speed perturbation paradigm in which 29 participants tracked a moving target that underwent brief speed changes. On each trial, we recorded eye velocity along with perceptual judgments of the sign of the speed change (faster or slower) and confidence ratings for those judgments. Across participants, perceptual and oculomotor sensitivity were correlated (r = 0.53), but trial-by-trial perceptual and oculomotor responses appeared independent. A logistic regression model incorporating efference copy gain to predict perceptual reports from stimulus speed and eye speed revealed systematic under-compensation for eye movements; faster eye movements were associated with lower changes in perceived speed. Confidence ratings were well-calibrated to perceptual accuracy and showed a consistent negative relationship with eye movement speed. Collectively, these results are consistent with systematic undercompensation for eye movements through the use of efference copy during motion perception.

## Introduction

The analysis of an object’s visual motion serves two primary purposes: motion perception and the control of eye movements. Motion perception refers to the brain’s ability to detect and interpret the changing position of objects in the visual field, while eye movement control ensures that visual inputs remain stable on the retina and behaviourally relevant (Pashler & Yantis, 2002; Schiller, 1998). Precise eye movement control is critical for supporting ongoing perception of the features of any object being tracked. Neural activity in dorsal stream areas including the middle temporal and medial superior temporal areas (MT/MST) has been closely associated with both perceptual judgements of motion and the generation of smooth pursuit tracking eye movements (Newsome et al., 1988). Despite this close association, the interdependence of visual motion perception and oculomotor control is poorly understood.

Two questions are of particular interest to us: first, to what degree are the same neural representations of motion information used to drive both perception and eye movements; and second, given that eye movements affect retinal motion, how does downstream perceptual and oculomotor processing compensate for the predicted effects of eye movements. One fruitful approach to investigating perceptual-motor interactions involves paradigms in which participants track objects with their eye movements and subsequently make perceptual judgements about the object’s velocity (speed or direction), or changes in velocity (Beutter & Stone, 2000; Gegenfurtner et al., 2003). Trial-by-trial correlations between the direction of eye movements and perceptual reports during smooth pursuit have been used to argue for shared neural mechanisms underlying the sensory processing contributing to the control of eye movements and perception (Beutter & Stone, 2000). Blum and Price (2014) extended this to reflexive ocular following, finding correlated directional biases across test directions consistent with shared sensory processing, but no trial-by-trial correlation between perceptual and oculomotor errors. Price and Blum (2014) further showed that across individuals, perceptual precision correlated with the precision of smooth pursuit but not ocular following, suggesting that volitional pursuit shares sensory neural constraints with perception while reflexive eye movements depend on partially independent circuits.

Speed processing, however, reveals a different pattern. Gegenfurtner et al. (2003) asked participants to track moving objects that underwent stepped changes in speed and to report the sign of the speed change. Despite finding matched speed discrimination thresholds for perception and pursuit, trial-by-trial analysis of pursuit errors and perceptual errors showed no correlation. Their finding indicates that pursuit and speed perception may depend on independent mechanisms, but it remains unclear how these systems can show matched sensitivity yet operate independently at the single trial level.

In the context of a closed loop feedback system for controlling smooth pursuit eye movements, the role of efference copy in modulating the sensory signals used to drive smooth pursuit is well characterised. Achieving visual stability during eye movements represents a fundamental computational challenge. However, eye movements also introduce motion into the visual information processed by the retina, creating retinal slip that causes motion blur (Lencer & Thrillenberg, 2008; Ilg, 2009). To counteract this, the brain integrates retinal signals with extraretinal signals such as efference copy, a duplicate of motor commands sent to sensory regions. Efference copy signals can be used to distinguish self-induced visual motion from external object motion and predict the sensory consequences of movements (Bridgeman, 1995, 2007; Ford & Mathalon, 2019). However, the degree to which efference copy perfectly compensates for the impact of eye movements on motion perception remains unclear.

Efference copy operates within a broader computational framework known as predictive coding, where the brain continuously generates predictions about expected sensory input and compares these predictions with actual sensory signals (Ford & Mathalon, 2019). During smooth pursuit, efference copy provides the specific signal that enables these predictions, effectively "subtracting" the sensory consequences of eye movements from retinal input, therefore stabilising perception. However, this system is not perfect. Systematic biases in speed adjustments could arise when there is a mismatch between predicted and actual sensory feedback. Such mismatches can result from neural transmission delays (∼50–100 ms), noisy sensory inputs, and rigid internal models that persist with outdated predictions. The direction of these biases (i.e. whether the brain over-estimates or under-estimates target speed), and their magnitudes remain incompletely quantified in the literature. Efference copy disruptions have been implicated in schizophrenia, where impaired smooth pursuit and motion perception may reflect dysfunctional predictive signals (Ford & Mathalon, 2019; Frith, 1987; Holzman et al., 1976; Silverstein, 2016; Spering et al., 2013).

To better understand the interactions between motion perception and oculomotor control, we designed a smooth pursuit paradigm in which participants tracked a moving target that underwent brief perturbations in speed. On each trial, we measured oculomotor responses (through eye tracking), perceptual judgments (through participants’ reports of whether they perceived speed changes as faster or slower than baseline), and perceptual confidence. First, we hypothesised that if the tracking and perceptual judgments depend on the same sensory representations, individuals with greater perceptual sensitivity should also display greater oculomotor sensitivity. Second, we hypothesised that if the perceptual system imperfectly compensates for the efference copy of the oculomotor command, human judgments of stimulus velocity during smooth pursuit should be biased by variations in eye velocity. Third, we predicted that participants’ confidence in their perceptual reports should be systematically related to efference copy mechanisms during smooth pursuit, evident as a correlation between the amplitude of error in tracking and confidence levels. We showed that perceptual and oculomotor sensitivity were correlated across individuals, but trial-by-trial responses appeared independent. To assess how effectively perceptual judgments remove the influence of eye movements on retinal velocity, we developed a logistic regression model to quantify efference copy compensation. This modelling suggests that the perceptual system systematically underweights the efference copy signal, compensating for only approximately 88% of the eye velocity contribution to retinal motion. Confidence ratings on each trial were closely related to perceptual accuracy and showed a consistent negative relationship with eye movement speed in the direction predicted by undercompensation. Collectively, our data suggest that imperfect efference copy during smooth pursuit is the source of a small but consistent perceptual bias whereby faster eye movements lead to underestimates of target speed.

## Methods

### Participants

Twenty-nine adult participants (22F, 7M) were tested in 2024-25, including volunteers who were compensated for their time and three authors. All participants had normal or corrected-to-normal vision and no known neurological or oculomotor conditions. The non-author volunteers had no, or very limited prior psychophysical testing experience and were naive with respect to the aims of the study. All procedures were approved by the Monash Human Research Ethics Committee, and volunteers gave informed written consent.

### Apparatus

Participants were comfortably seated facing a computer monitor (Display++; Cambridge Research Systems Ltd., 120 Hz refresh rate; 1920 × 1080 pixels; 705 × 395 mm) with a chin and forehead rest used to stabilize the head and maintain a viewing distance of 570 mm. During the experiment, participants were asked to limit head movements and only move their eyes. Eye movements were recorded using a non-invasive, video-based eye-tracker with a sampling frequency of 1000 Hz (Eyelink 1000, SR Research, Ottawa, Canada). The system has a spatial resolution of 0.01° and average accuracy of 0.25-0.5°. Stimuli were generated using MATLAB R2024a (The MathWorks, Natick, MA) and the Psychophysics Toolbox extensions (Kleiner et al., 2007) under the control of custom code in Neurostim (https://github.com/klabhub/neurostim).

### Stimuli and Task

A stationary white target (0.5° diameter, 234 cd/m²) appeared against a neutral grey background (117 cd/m²). After the participant acquired fixation of the target, it started moving at a constant velocity of 4°/s. After 700 or 750 ms, the target changed speed for 200 or 100 ms, then returned to its original speed until the end of the trial. The period of motion was therefore always 1600 ms, with the speed perturbation centred within the trial. These brief perturbation durations were selected to minimise within-trial adaptation (Madelain & Krauzlis, 2003), allowing us to capture immediate perceptual-motor interactions before recalibration can occur.

Participants 1-13 completed trials with a 100 ms speed perturbation, with speed changes of [-2.5, -1, -0.5, -0.25, 0, 0.25, 0.5, 1, 2.5°/s] relative to the baseline 4°/s. Participants 14-29 completed trials with a 200 ms perturbation and changes of [-2.5, -1.5, -0.75, -0.25, 0, 0.25, 0.75, 1.5, 2.5°/s]. The 200 ms condition was introduced after the 100 ms speed changes proved difficult to judge.

Participants were asked to follow the moving target as closely as possible. At the end of the trial, they were asked to report whether they perceived an increase or decrease in speed by pressing the Up or Down arrow on the keyboard. Participants were also instructed to double-click the same button to indicate high confidence about their decision.

### Procedure

Our study design was based on previous work by Gegenfurtner et al. (2003), who used speed changes lasting 200 and 1000 ms. We employed shorter perturbation durations (100 and 200 ms) to examine perceptual decisions during the early, transient phases of pursuit adaptation, when efference copy compensation might be most critical and potentially imperfect.

Before each block of trials, participants completed an eye tracking calibration procedure requiring them to fixate targets in a 9-point grid. The calibration was accepted if the average error was less than 0.5° and no individual point had an error exceeding 1°. The experiment consisted of four blocks of trials, with each containing 72 trials (8 repeats of 9 speed changes, in randomized order). Each trial was followed by auditory feedback indicating whether their perceptual judgment was correct or incorrect. Trials with eye position errors exceeding 4 degrees or interrupted by blinks were immediately terminated and reshown later in the random sequence.

### Psychometric Analysis

We fit psychometric responses for each participant (proportion of “faster” responses at each speed change), with logistic functions using maximum likelihood methods implemented in the MATLAB Palamedes toolbox. Goodness of fit was assessed using R² values, with a threshold of R² ≥ 0.7 for psychometric fits considered acceptable. From these fits, we extracted the Point of Subjective Equality (PSE) and sensitivity (slope) parameters. The PSE represents the participant’s bias, or the speed change magnitude at which perceptual judgments were equally likely; p(Faster)=50%. For an unbiased observer, the PSE should be close to 0. Sensitivity (β) is defined as the slope of the fitted logistic function evaluated at the PSE, steeper slopes correspond to higher sensitivity to speed changes.

### Oculometric analysis

Eye position data sampled were recalibrated, smoothed, and desaccaded before extracting metrics to summarize performance. Eye positions on each trial were recalibrated to compensate for slight head movements between trials that artificially inflate errors in eye position. The recalibration process allowed horizontal and vertical translation to minimise eye position errors during the fixation period at the start of the trial. The recalibrated eye position data were smoothed and differentiated using a Savitzky-Golay filter with order 3 and a frame length of 51 samples. Saccades were detected when horizontal or vertical eye speed exceeded 20°/s. Data from 15 ms before to 15 ms after each saccade were replaced with NaN and excluded from subsequent analyses, allowing the remainder of the trial to contribute to trial averaging.

To transform continuous eye velocity measurements into a format comparable to binary perceptual judgments, we adopted an oculometric analysis based on the Receiver Operating Characteristic (ROC). For each speed change magnitude, we compared the distribution of eye velocities on each trial to the distribution when no speed change occurred (0% speed change magnitude). The area under the ROC curve (AUC) represents the discriminability between conditions, with values ranging from 0.5 (chance-level discrimination) to 1.0 (perfect discrimination). AUC were calculated separately for each speed change to form an oculometric function, which we fit with logistic functions in the same manner as the psychometric functions. Due to the inter-trial variability of eye movements, we accepted fits with R² ≥ 0.6.

To examine how oculometric sensitivity changed across time, we examined 100 ms wide windows in 50 ms steps from 0–400 ms after the speed change. The 150–250 ms window yielded highest oculometric sensitivity and is used for most analyses, but the precise choice of window did not qualitatively affect our results.

### Data Inclusion Criteria

A participant’s data was included in each analysis only if they met the following quality criteria. Psychometric functions were accepted if there were ≥5 trials per speed change condition, the model converged, the sensitivity was positive and R² ≥ 0.70. Oculometric functions were accepted using the same trial count and model convergence criteria, with the additional requirement that AUC values showed a significant monotonic relationship with speed change (Spearman ρ > 0.5, p < 0.05). Of the 29 participants tested, 22 met psychometric criteria (100 ms: 8/13; 200 ms: 14/16) and 28 met oculometric criteria (100 ms: 13/13; 200 ms: 15/16); 21 met both (100 ms: 8/13; 200 ms: 13/16). One participant of the 21 who met both psychometric and oculometric criteria was excluded from the efference copy modelling because of poor model fits.

### Efference Copy Gain Model

We estimated the gain of efference copy using a model that assumes that the brain has access to a reliable estimate of the object’s retinal slip speed but imperfectly compensates for efference copy signals of the eye velocity to determine perceived velocity. Therefore:

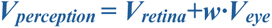

where w represents the efference copy gain parameter. Perfect compensation occurs when w=1, while w=0 indicates no compensation. As the retinal velocity is the difference between object and eye velocity:

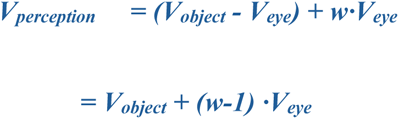

In our experiments, V_object_ is manipulated and V_eye_ and V_perception_ are measurable, allowing us to determine the efference copy gain w. We implemented this within a logistic regression framework modelling whether a participant reports a speed change as “faster” than baseline:

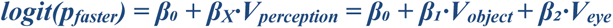

In Equation 3, β_0_ represents the baseline bias (the intercept term that determines the PSE when both V_object_ and V_eye_ are zero). β_x_ represents the perceptual sensitivity to speed changes. β₁ is the slope parameter representing the sensitivity to object velocity; β₂ is the slope parameter for eye velocity, which represents the uncompensated portion of eye movements that biases perception.

By fitting this model to each participant’s data, we estimated both their efference copy gain (w) and the corresponding eye velocity coefficient (β₂).

To assess whether eye velocity contributed significantly to perceptual predictions at the individual level, we compared the full model (β₀ + β₁·V_object_ + β₂·V_eye_) to a reduced model (β₀ + β₁·V_object_) using a likelihood ratio test (chi-square, df = 1) and Akaike Information Criterion (AIC). A participant was considered to show a significant eye velocity effect if the likelihood ratio test reached p < 0.05 or ΔAIC < 0. At the group level, we tested whether the mean β₂ differed significantly from zero using a one-sample t-test.

### Quantifying meta-cognition

On each trial, confidence was recorded as either low or high: a single press on the keyboard indicated low confidence and a double-click indicated high confidence. Participants who used only one confidence level were excluded from metacognitive analyses (n=3). As a formal measure of metacognitive sensitivity, we estimated meta-d’ and the M-ratio (meta-d’/d’) using the method of Maniscalco and Lau (2012), implemented using fit_meta_d_MLE. Trials were pooled across all non-zero speed change magnitudes, with responses recoded so that a correct response was consistently labelled as the signal category (n=26).

To examine whether perceptual confidence relates to eye movement strength — an analysis that, to our knowledge, has not previously been reported — we computed a z-scored eye speed difference (z_diff_). For each non-zero speed change magnitude, trial-level eye speeds were z-scored and outliers beyond two standard deviations removed and re-z-scored. Z_diff_ was the difference in mean z-scored eye speed between high- and low-confidence trials, sign-inverted for decrement magnitudes so that the sign of z_diff_ is consistent across increment and decrement trials. Magnitudes with fewer than five trials in either confidence bin were excluded; participants required at least three valid magnitudes (n=24, including participants where the efference copy model was attempted but did not meet all quality criteria for gain estimation). A one-sample t-test was used assess if the group mean z_diff_ differed from zero.

## Results

We compared perceptual judgements and oculomotor sensitivity to speed changes during constant-velocity smooth pursuit. Below, we first explore inter-individual differences in psychometric and oculomotor sensitivity, then examine if psychometric and oculomotor performance are correlated at the single trial level. Finally, we develop a model of speed perception during eye movements that incorporates efference copy and use it to infer how the gain of efference copy influences motion perception.

### Oculomotor and psychometric sensitivity are correlated

We obtained a perceptual report and smooth pursuit eye tracking associated with speed changes on each trial (Figure 1A). We first separately examined psychometric and oculomotor performance. Psychometric performance was quantified using the proportion of trials at each speed change reported as containing an increment, p(Faster), and fit with a logistic regression function (Figure 1B). We focus on the point of subjective equality (PSE) and sensitivity (β). Pursuit eye movements systematically deviated from the baseline 4 deg/s during and after the speed perturbation (Figure 1C), and we quantified the corresponding oculomotor performance using an area under the ROC curve approach (AUC). For each stimulus condition, the AUC compares the distribution of eye speeds across trials with the distribution of eye speeds associated with the no-change condition (Figure 1D).

**Figure 1.**
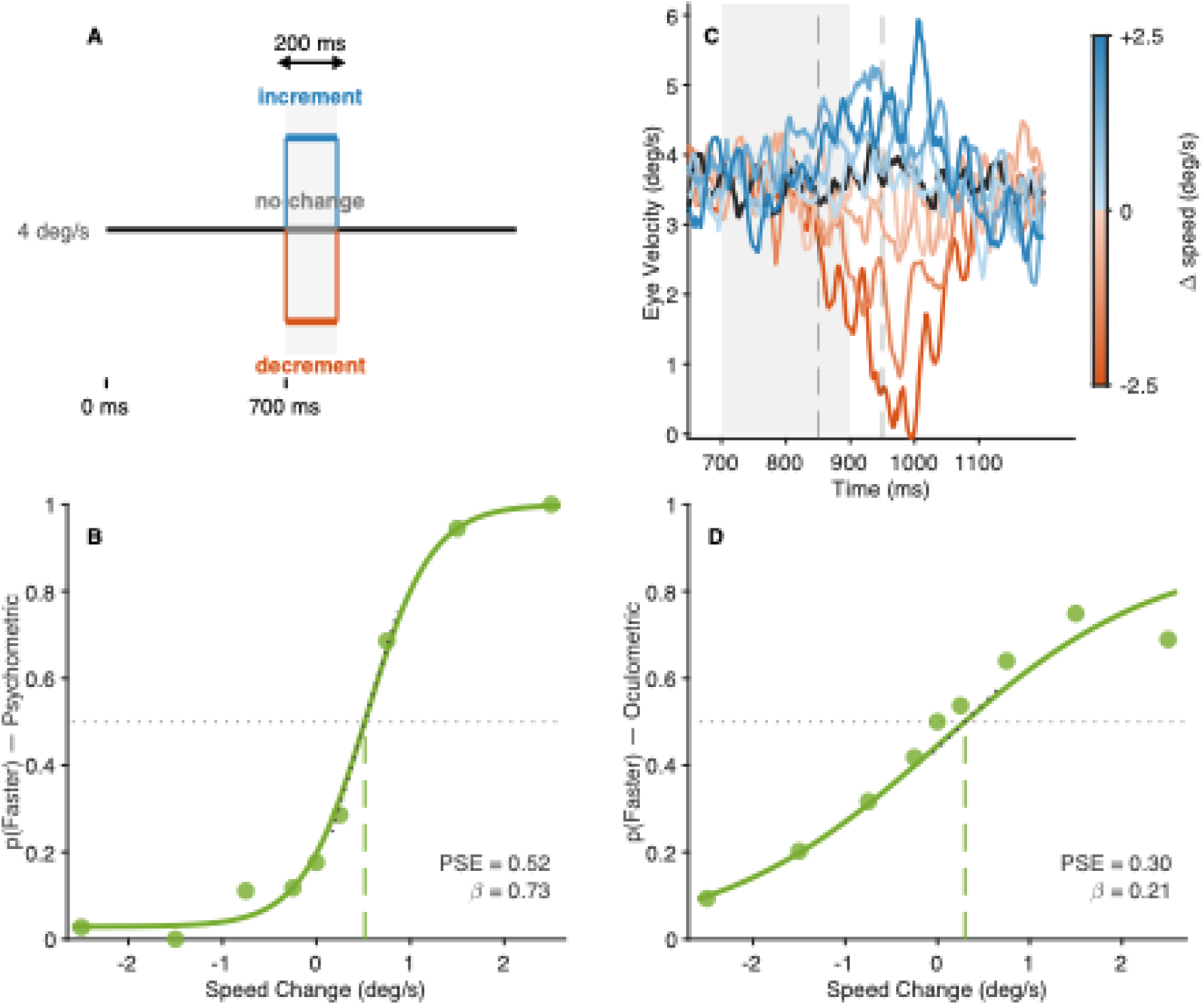
Task design and example psychophysical and oculomotor performance for a single participant. (A) Participants tracked a target moving at 4 deg/s that underwent a 200 ms speed change beginning 700 ms after trial onset. Three perturbation types are shown: increment (blue), no change (black), and decrement (orange). For participants completing trials with the 100 ms perturbation condition, the structure was identical with perturbation onset at 750 ms. (B) Psychometric function showing the proportion of trials judged "faster" as a function of speed change magnitude. Filled circles represent trial-averaged proportions; solid curve shows the fitted logistic function. The vertical dashed line marks the point of subjective equality (PSE) and the dotted line indicates the slope at the PSE from which sensitivity (β) was derived. (C) Trial-averaged eye velocity traces colour-coded by speed change magnitude. The grey shaded region indicates the perturbation window; dashed vertical lines mark the analysis window 150–250 ms after perturbation onset used for most oculometric analyses. Line colour indicates speed change (orange, decrements; blue, increments; darker colours, larger magnitudes); the no-change trace is black. (D) Oculometric function derived from receiver operating characteristic (ROC) analysis. Filled circles show the area under the ROC curve (AUC) for each speed change magnitude. The dashed vertical line marks the oculomotor PSE and the dotted line indicates the slope at the PSE from which oculometric sensitivity (β) was derived.

In Figure 1C it is clear that changes in eye speed were delayed and smoothed with respect to the changes in stimulus speed. To determine the eye velocity analysis window that optimised our ability to quantify oculometric sensitivity, we examined how two metrics varied time after perturbation onset. First, we determined the slope of the AUC–speed change relationship (Figure 2A), which peaked at 200 ms, i.e. eye speeds from 150-250 ms after speed change onset. Second, we calculated the Spearman correlation between the AUC and speed change magnitude. This measure of consistency also peaked at 200 ms (Figure 2B). Therefore, all subsequent oculometric analyses focused on this 150-250 ms time window, however, the exact choice of time window did not qualitatively affect the interpretation of our results.

**Figure 2.**
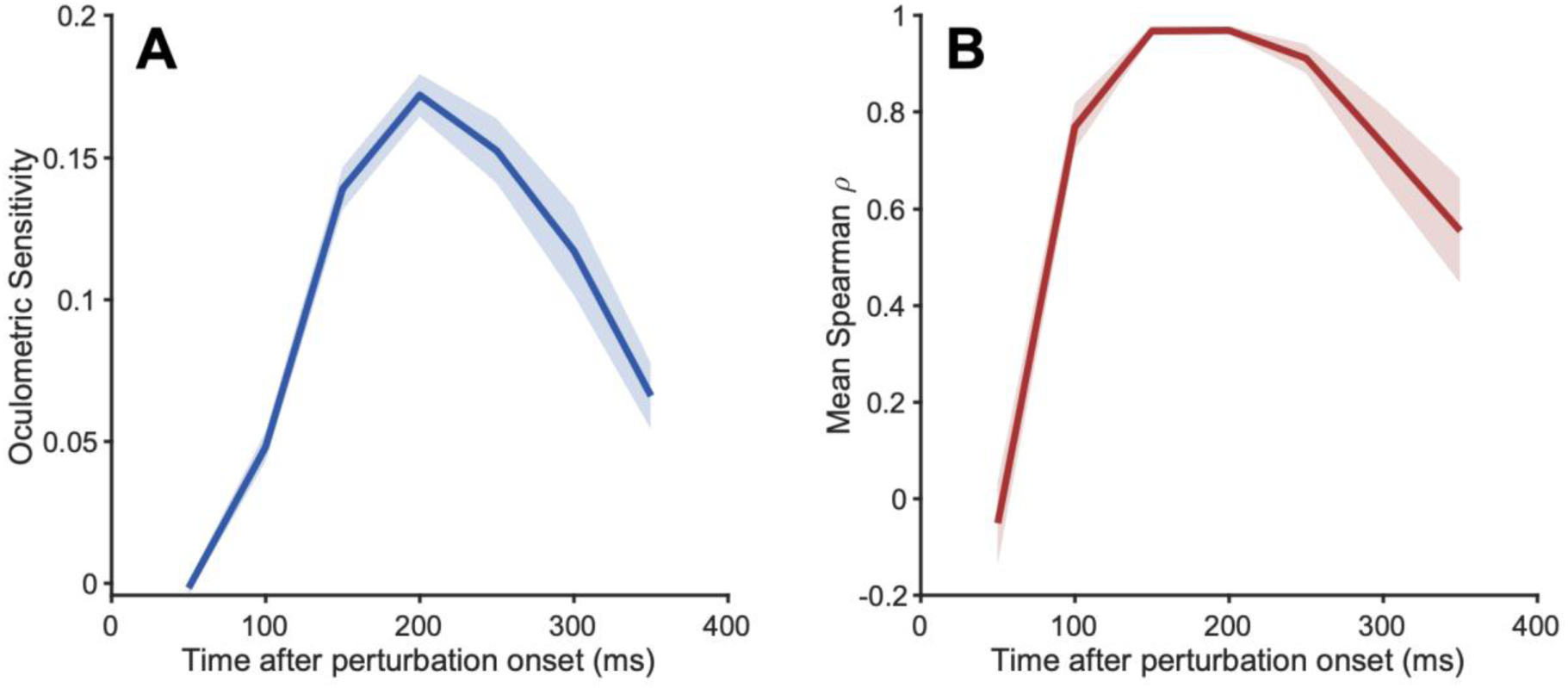
The 150–250 ms post-perturbation window maximises oculometric sensitivity. Oculometric sensitivity (A) and Spearman correlation between changes in stimulus and eye speed (B) were calculated in 100 ms windows with centres from 50-350 ms after perturbation onset. Mean (SEM) across 21 participants are shown.

If motion perception and pursuit depend on shared neural mechanisms, metrics of perceptual judgments and oculomotor tracking should be correlated in some way across participants. Psychometric and oculometric functions were fitted for each participant, and participants were included only if both fits met quality criteria (see Methods). We examined whether individual differences in PSE and sensitivity were correlated across all participants, and separately for speed changes of each duration (Figure 3). Mean PSEs showed no significant differences between perceptual and oculomotor systems for any grouping, and we observed no cross-system correlation for any grouping (Figure 3A-C. t-test and Pearson correlation, all p>0.1). Critically, oculometric and perceptual sensitivities were correlated across participants (Figure 3D-F). The correlation was not significant in the 100 ms condition alone (r = 0.46, p = 0.253), but was significant in the 200 ms condition (r = 0.57, p = 0.042) and across all participants (r = 0.53, p = 0.014). Note that when calculating the correlation for all participants, sensitivities were first z-scored separately within each duration condition before combining. Perceptual and oculometric sensitivity were not significantly different from one another in either condition (100 ms: t(7) = −1.35, p = 0.214; 200 ms: t(12) = −0.88, p = 0.394), consistent with previous reports of matched thresholds between the two systems (Beutter & Stone, 2000; Gegenfurtner et al., 2003). We note that as estimates of oculometric sensitivity depend heavily on the choice of analysis time window, this consistency should be interpreted with caution.

**Figure 3.**
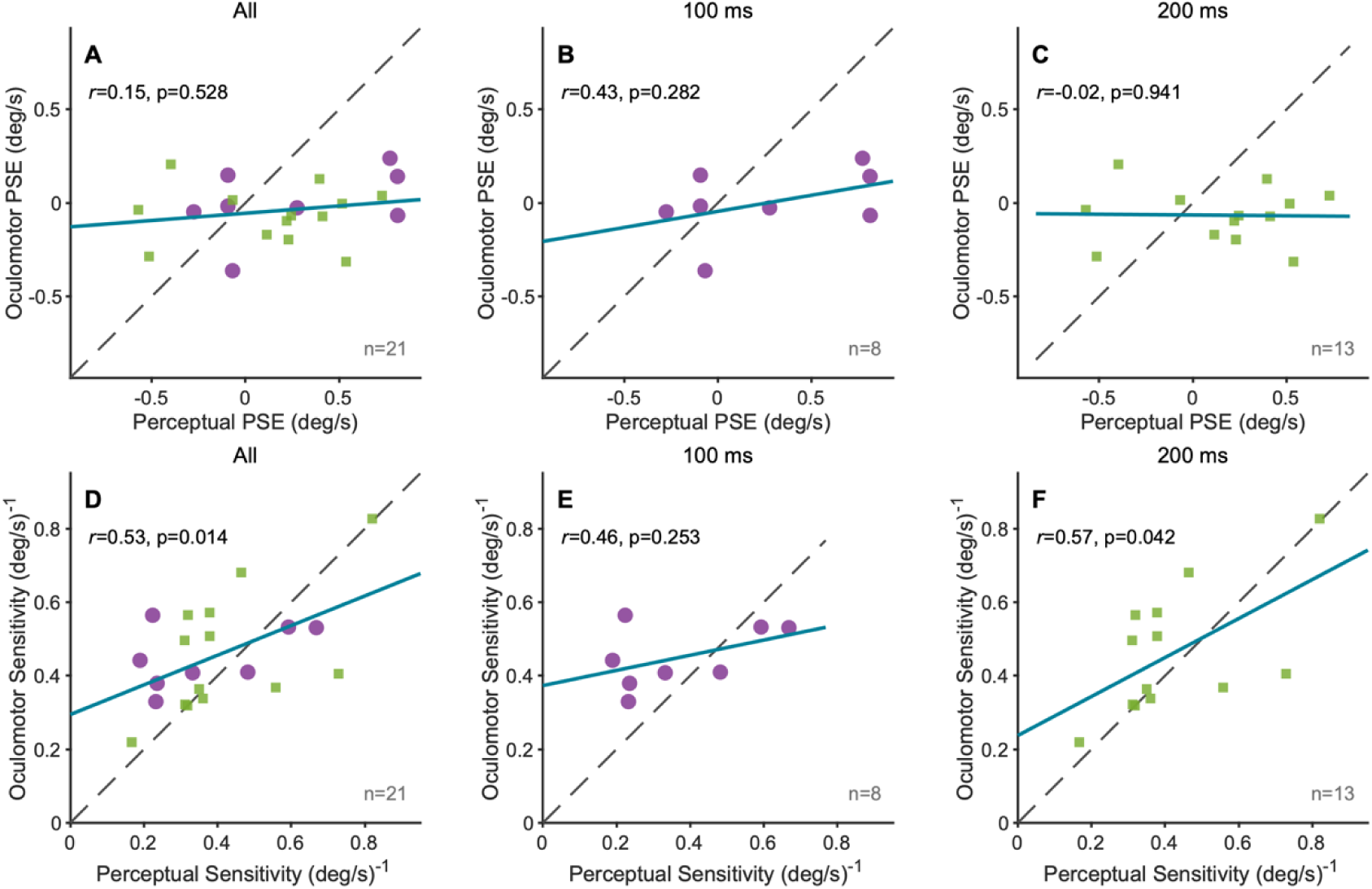
Perceptual and oculomotor sensitivity, but not PSEs, are correlated across participants. PSEs (A-C) and Sensitivity β (D-F) are compared for oculometric and perceptual reports. Data are shown from all participants collapsed across speed change durations (A, D), and separately for participants who completed the 100 ms condition (B, E) or 200 ms condition (C, F). Each dot represents one participant.

### Perceptual judgments and eye velocity fluctuations appear uncorrelated on individual trials

Given the correlated sensitivity observed across participants, we wondered how an individual’s fluctuations in perception and eye velocity might co-vary across trials. Thus, we assessed the trial-by-trial correspondence between perceptual responses and eye movements using the proportion of trials with matched outcomes (p_Match_) between perceptual reports ("faster" or "slower”) and oculomotor responses (eye speed faster or slower than the median across all trials and all conditions) (Beutter & Stone, 2000; Stone & Krauzlis, 2003). Under the null hypothesis of independence, p_Match_ should equal the chance level (p_Chance_), calculated separately for each speed change magnitude as:

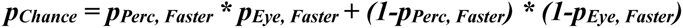

where p_Perc, Faster_ and p_Eye, Faster_ are the proportions of "faster" responses from the psychometric and oculometric functions, respectively. This formula accounts for both types of matches: trials where both the perceptual report and change in eye speed were classified as "faster" (p_Perc_, _Faster_ * p_Eye, Faster_) and trials where both were classified as "slower" ((1-p_Perc, Faster_) * (1-p_Eye, Faster_)). Neither individual participants (binomial tests, all p > 0.05) nor the group showed significant deviations from chance at any speed change magnitude (one-sample t-tests, all p > 0.05). At speed change magnitude = 0, p_Match_ remained near 50% (Figure 4), consistent with independent processing. The U-shaped increase in p_Match_ at larger absolute speed change magnitudes is consistent with both systems having similar sensitivity to stimulus strength.

**Figure 4.**
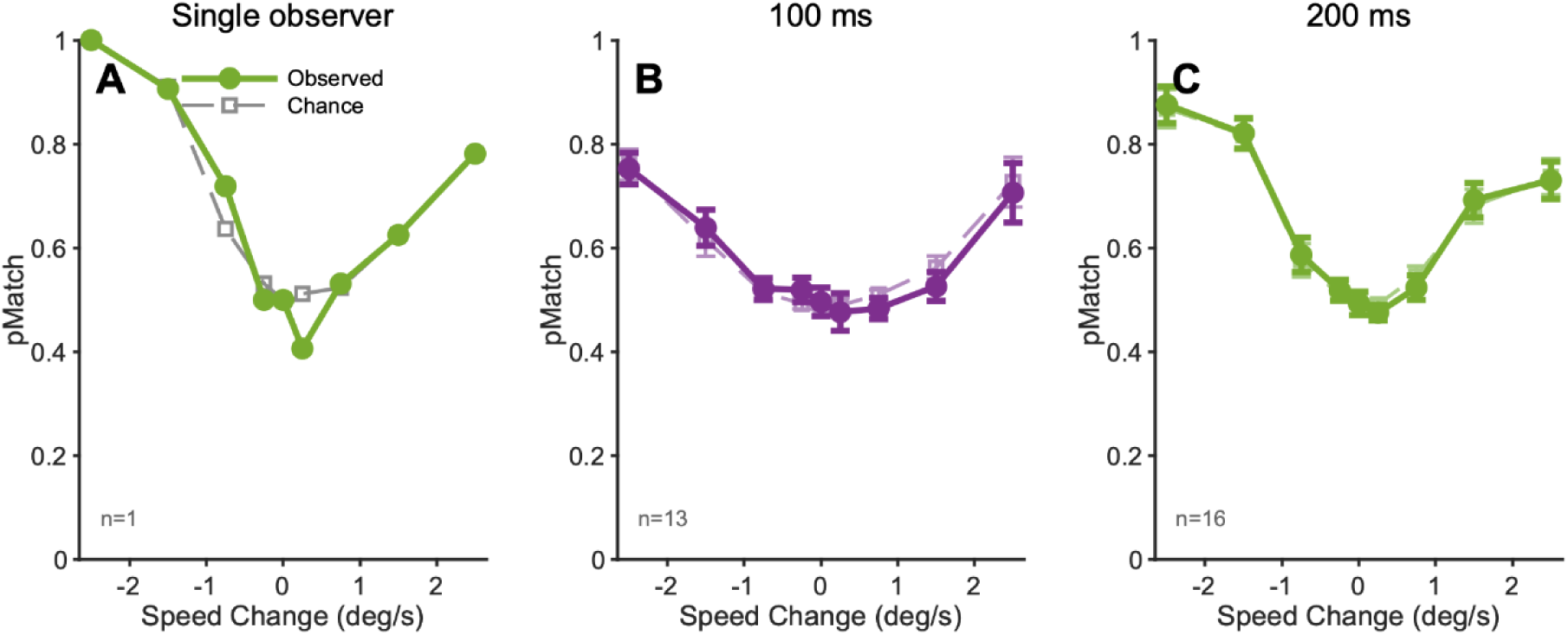
Perceptual judgments and eye velocity fluctuations are uncorrelated on individual trials. The proportion of trials with matched outcomes (p_Match_) between perceptual decisions and oculomotor responses is plotted as a function of speed change magnitude. p_Match_ was compared to the theoretical chance level (p_Chance_) expected under independence, calculated from the psychometric and oculometric functions for each speed change magnitude. (A) Example participant from the 200 ms condition. Group mean p_Match_ and p_Chance_ for the 100 ms (B; n = 13) and 200 ms (C; n = 16) conditions. Error bars indicate 1 SEM. No significant deviations from chance were observed at any speed change magnitude in either condition (one-sample t-tests, all p > 0.05). The U-shaped increase in p_Match_ at larger magnitudes reflects both systems showing similar sensitivity to stimulus strength.

### Efference copy compensation is systematically imperfect

Despite establishing that trial-by-trial perceptual and oculomotor responses appear independent, we next asked whether eye velocity nonetheless influences perceptual reports in a systematic way that can be explained by imperfect efference copy compensation. We developed a model to predict perceptual speed judgments assuming that the brain has access to a reliable estimate of the object’s retinal slip speed but imperfectly compensates for efference copy signals of the eye velocity (see Methods). We used this efference copy gain model to quantify how accurately the perceptual system compensates for ongoing eye movements during smooth pursuit (Figure 5). If compensation were perfect, the gain parameter *w* would equal 1; values below 1 indicate systematic undercompensation, meaning that on trials with faster eye movements, perception will be biased toward "slower" judgments. Across all participants with good model fits (n=20, one participant excluded due to a poor model fit), the mean efference copy gain (w = 0.88 ± 0.04) was significantly below 1.0 (one-sample *t*-test, *t*(19) = −2.78, *p* = 0.012, *d* = −0.62). While the same trends were seen when separately considering the participants tested with each speed change duration, neither perturbation condition had w significantly different from 1 (100 ms: t(6) = -1.92, p = 0.103, d = -0.73; 200 ms: t(12) = -1.96, p = 0.073, d = -0.54), and the two conditions did not differ from each other (t(18) = -0.87, p = 0.396). Efference copy gain was not significantly correlated with perceptual sensitivity across participants (r = −0.10, p = 0.669), consistent with the precision of speed discrimination and the degree of efference copy compensation being partially independent.

**Figure 5.**
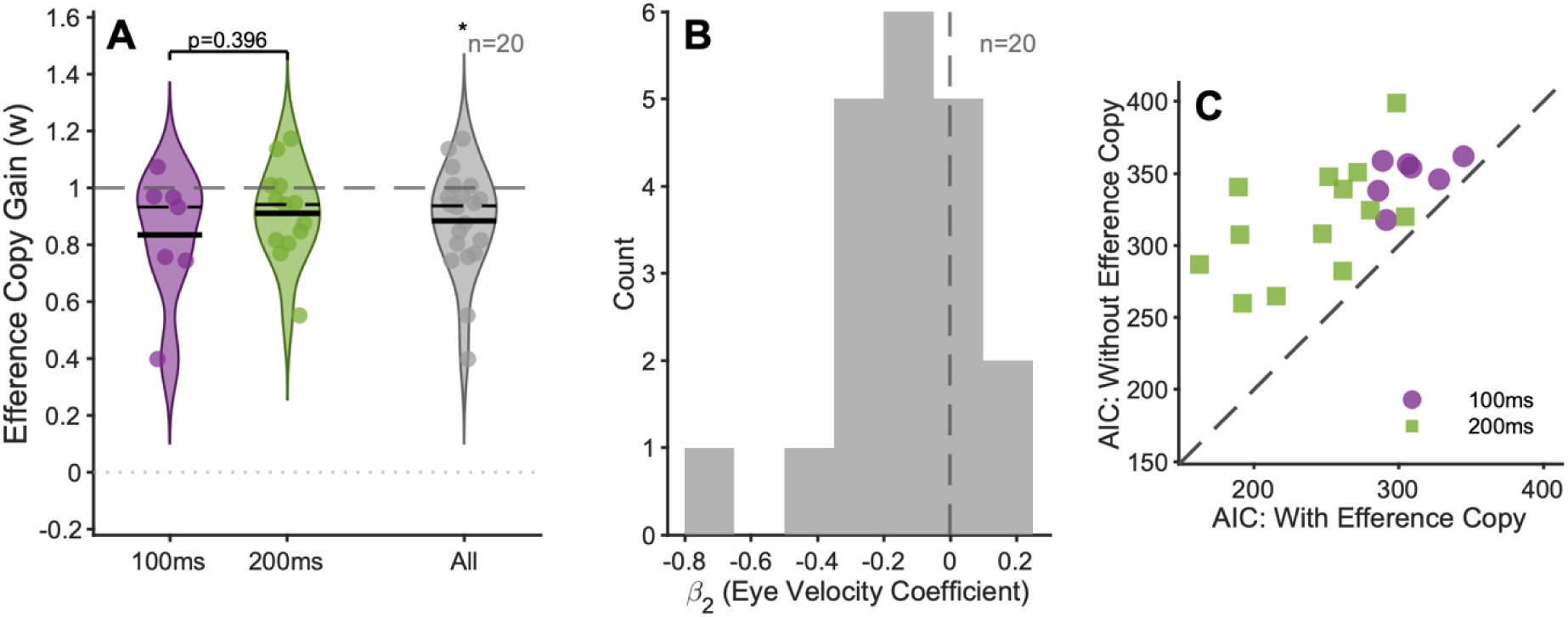
The perceptual system imperfectly compensates for eye movement signals. (A) Distribution of efference copy gain (w) derived from a logistic regression model for the 100 ms condition (purple), 200 ms condition (green), and all participants combined (grey), displayed as violin plots with individual data points overlaid. Solid and dashed lines show group mean and median. (B) Distribution of the eye velocity regression coefficient (β₂) across participants. Values below zero indicate that faster eye movements biased perception toward "slower" judgments. (C) Model fit comparison using Akaike Information Criterion for a model with Efference copy, including retinal slip and eye velocity, (x-axis) and a retinal slip-only model without Efference copy (y-axis). Points above the diagonal indicate that the Efference copy model provided a better fit.

We further characterised the robustness of the eye velocity effect in two ways. First, we examined β₂, which quantifies the influence of eye velocity on the perceptual report (Figure 5B). At the group level, mean β₂ was significantly negative (β₂ = −0.14, t(19)=-3.010, p=0.007), confirming a systematic eye velocity bias across participants. We separately used the Akaike Information Criterion (AIC) to compare the full model (incorporating V_eye_ and V_retina_) to a reduced model excluding V_eye_ (Figure 5C). The full model provided substantially better fit, with ΔAIC > 0 for all 20 participants and likelihood ratio test (all p < 0.05) (Figure 5C). This indicates that eye velocity information contributes statistically significant predictive power about perceptual reports beyond retinal slip alone.

### Confidence predicts perceptual accuracy

To assess metacognitive awareness, we first examined whether participants’ confidence ratings meaningfully reflected their perceptual performance (Figure 6). This validation step is essential, as any interpretation of confidence-related effects requires establishing that our confidence measure captures genuine metacognitive processes rather than response biases.

**Figure 6.**
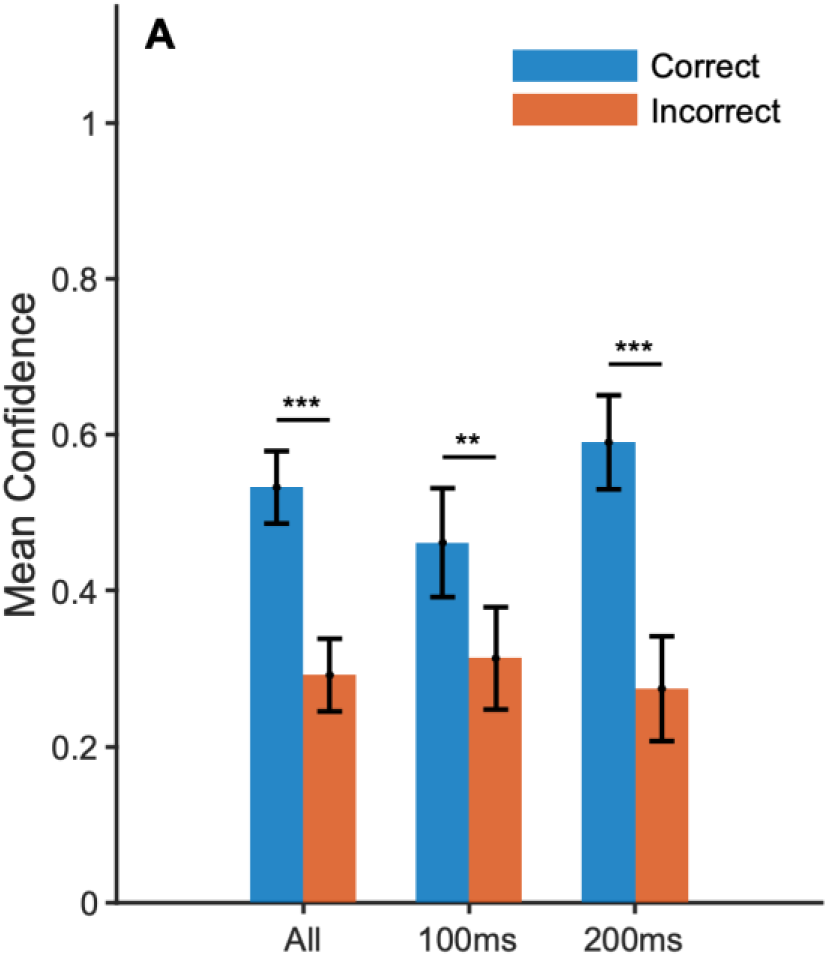
Confidence ratings reflect perceptual performance. Grouped bar charts show mean confidence values for correct and incorrect responses for all participants combined (All), the 100 ms condition, and the 200 ms condition separately. Error bars indicate ±1 standard error of the mean (SEM). Significance brackets indicate the outcome of paired t-tests comparing the two bar types within each group: *** p < 0.001, ** p < 0.01. Participants expressed significantly higher confidence on trials they answered correctly than on trials they answered incorrectly, across all participant groups.

Across all participants (n=29), confidence was strongly related to accuracy. Participants expressed significantly higher confidence for correct trials (mean confidence = 0.53 ± 0.05 SEM) than incorrect trials (mean confidence = 0.29 ± 0.05 SEM; paired t-test, t(28)=6.66, p<0.001, d=0.87) (Figure 6). Perceptual metacognitive efficiency estimated using standard meta-d’ (Maniscalco & Lau, 2012) yielded a mean M-ratio of 0.93 ± 0.23, consistent with ideal efficiency (t(25) = −0.29, p = 0.77). We also examined whether confidence predicted eye movement speed using z_diff_. A negative z_diff_ indicates that eye speeds were relatively lower on high-confidence than low-confidence trials. The mean z_diff_ (quantifying whether eye movement speed predicted confidence) showed a consistent negative trend (mean = −0.048 ± 0.024, t(23) = −1.99, p = 0.059), consistent with systematic undercompensation biasing high-confidence trials toward slower eye movements.

## Discussion

We measured perceptual and oculomotor responses to brief speed perturbations during smooth pursuit. Our two main findings were: (1) perceptual and oculomotor sensitivity were correlated across individuals, with participants who showed higher oculomotor sensitivity also tending to show higher perceptual sensitivity; and (2) mean efference copy gain was significantly less than 1, indicating that participants systematically undercompensated for their own eye movements when perceptually reporting speed changes. Below, we discuss how these findings give insights into common sources of sensory processing underlying perception and oculomotor control, and how eye movements directly and indirectly influence motion perception.

### Common sensory sources for perception and oculomotor control

If motion perception and the control of tracking eye movements depend on the same networks of sensory neurons, we should expect to observe correlations between perceptual motion sensitivity and the accuracy, precision or gain of eye movements. These correlations could be present *across trials* or *across individuals;* we examine these possibilities separately below.

At the level of *trials*, trial-by-trial correlations have commonly been reported for oculometric and psychometric direction judgments (Beutter and Stone, 2000; Krauzlis and Adler, 2001; Stone and Krauzlis, 2003), but not for tasks requiring speed, or speed-change, judgements during pursuit (Gegenfurtner et al., 2003; Tavassoli and Ringach, 2010), or during ocular following (Bostrom and Warzecha, 2010; Spering et al., 2011; Blum and Price, 2014; Glasser and Tadin, 2014). Our modelling showed that most observers had an efference copy gain (w) less than 1, implying that fluctuations in eye velocity introduce consistent biases into perceptual decisions, shifting the psychometric function. This systematic bias should also generate trial-by-trial covariance between perceptual and oculomotor systems. Therefore, if this model is valid, it suggests that independent noise sources sufficiently reduce the covariance to levels that cannot be reliably measured with the number of trials our participants completed.

If the oculomotor and perceptual systems share common noise sources, it is difficult to reconcile both the apparent absence of correlations in tasks involving speed judgements, and the difference in results between tasks involving direction versus speed judgements. This may reflect different stimulus paradigms, as many of the direction-judgment tasks referenced above used long duration stimuli, commonly 1000 ms. In contrast, all of the studies using ocular following used much shorter durations, and most pursuit studies, including ours, use shorter tracking durations (typically 200 ms, although Gegenfurtner et al also used a 1000 ms duration). Another critical difference is that tasks assessing speed sensitivity primarily focus on responses to *changes* in stimulus speed or the impact of stimulus onset, whereas tasks assessing direction sensitivity allow observers to integrate information throughout each entire trial.

In order to understand why the coupling between perception and eye movements appears to depend on the specific motion feature being processed, it would be useful to assess sensitivity to changes in speed, direction, or velocity (i.e. both) in the context of a common task design.

Relatively few studies have compared sensitivity across *individuals*, in part because it depends on large sample sizes. Note that we do not attempt to directly compare perceptual and oculomotor sensitivity, and such comparisons should be interpreted cautiously. Oculomotor sensitivity is a derived measure that depends on how neural activity is decoded, and the time windows that inform perception and eye movements may be substantially different. Influences such as attention and vigilance may also differently affect perception and eye movements. However, correlations between perceptual and oculomotor sensitivity do offer insights. We have previously observed correlated “within-domain” performance between two perceptual tasks assessing the precision of direction judgements and separately between two oculomotor tasks assessing the precision of eye movement direction (Price & Blum, 2014). In the same study, we also observed correlated “between-domain” performance between perceptual and oculomotor direction sensitivity during a pursuit task. This between-domain correlation was replicated here, with perceptual and oculomotor sensitivity, but not PSEs, significantly correlated across participants. Recent studies with large participant numbers also support the concept of an “oculomotor signature”, with robust individual differences between participants, that show correlations between motion sensitivity and oculomotor control (Bargary et al. 2017, Yan, Wu…Price et al. 2025, Goettker and Gegenfurtner, 2025).

### Influences of eye movements on perception

How do eye movements influence perception? The previous section explored how correlated oculomotor and perceptual sensitivity might reflect common sensory representations underlying oculomotor and perceptual responses to the same visual motion. However, these correlations might arise if eye movements influence motion perception. This could happen directly, due to perception of the retinal slip generated by eye movements, or indirectly, due to an efferent copy of the motor command targeting the sensory circuits underlying perception.

Our efference copy model provides a direct test of this distinction. A perceptual system relying on retinal slip alone should be well described by a model using this term without any additional contribution from eye velocity itself. However, the coefficient β₂ on V_eye_ isolates the indirect route, since it captures any influence of eye velocity on perception beyond what retinal slip already predicts. A significant, non-zero β₂ is therefore direct evidence that efference copy, rather than retinal slip alone, shapes perceptual reports during pursuit. We found β₂ was significantly negative at the group level, and a model including both retinal slip and eye velocity outperformed a retinal-slip-only model for all 20 quality-filtered participants. Thus, we conclude that perception during smooth pursuit is not a pure readout of retinal motion; it is shaped by an efference copy signal that only partially cancels the eye’s own contribution to that motion. If this compensation is incomplete rather than simply noisy, the same imperfect signal that biases the perceptual decision might also enter the confidence judgment, implying that confidence is not computed from a separately more accurate readout of eye velocity, but shares the decision’s own imperfect evidence. The negative z_Diff_ trend we observed is consistent with this possibility, though the effect was underpowered to confirm it directly.

### What can we learn from inter-individual differences?

We know that perception and pursuit likely share sensory representations of motion, given individuals with more precise perceptual sensitivity also show more precise oculomotor sensitivity. However, if efference copy links perceptual and motor systems by compensating for self-generated retinal motion, do individuals with more complete compensation also show higher perceptual sensitivity, or is compensation a separate sensorimotor factor? Comparing the fidelity of the extraretinal signal and the precision of early sensory processing across individuals may give insights into whether there are generalised neural mechanisms for signal representation throughout sensory and motor cortex. While our sample size was limited, in our sample, efference copy gain was not correlated with perceptual sensitivity, suggesting that compensation accuracy reflects a distinct factor. Future studies could address this more directly by quantifying efference copy gain and visual motion sensitivity in independent tasks, and using large cohorts suited to studying inter-individual differences.

Beyond its value for understanding healthy variation, quantifying efference copy gain may also have translational relevance. If gain varies continuously across the healthy population, clinical populations may either occupy the extremes of the distribution, or belong to a distinguishable group. For example, people with schizophrenia have impaired smooth pursuit gain; independently, they have disrupted predictive signalling, and saccadic control, reflecting a limited ability to translate motor commands into usable efference copy signals (Spering et al. 2013; Thakkar & Rolfs, 2019).

Positive symptoms of schizophrenia, such as auditory hallucinations and passivity experiences, are thought to arise from a failure of corollary discharge to distinguish self-generated events from those that are externally generated (Feinberg, 1978; Frith, 1987). Patients with hallucinations and passivity experiences fail to attenuate the perceived intensity of self-produced sensory stimuli relative to the controls (Blakemore et al.,2000), and corollary discharge dysfunction might explain auditory hallucinations, in which neural responses to self-generated speed are insufficiently suppressed (Ford & Mathalon, 2005; Ford & Mathalon, 2019). Corollary discharge failure has also been implicated in delusional ideation in healthy non-clinical samples (Malassis, Del Cul, & Collins, 2015). We predict that the general limitation in translating motor commands into usable predictive signals in people with schizophrenia or suffering hallucinations or delusions will manifest as lower low efference copy gain quantified using our approach.

Three limitations of the current study deserve consideration. First, our sample sizes after quality filtering were modest, with just 20 participants meeting our inclusion criteria for the efference copy model. Second, we collapsed across two speed-change durations. Third, the binary confidence measure, while sufficient to establish metacognitive validity and estimate perceptual M-ratio, limited statistical power for meta-d’ estimation given the relatively small number of trials per confidence bin. Given the challenges we faced with the short stimulus duration, future studies should focus on a longer 200 ms duration and would need to employ a confidence scale with more grades and higher trial counts to enable better metacognitive efficiency estimates. Extending the paradigm to clinical populations, particularly those with schizophrenia, would test whether the efference copy gain parameter provides a quantitative marker of predictive signal disruption, and whether it correlates with specific symptom dimensions.

## Acknowledgements

This project was funded by the Australian Research Council (DP200100179; DP210102107). MAH was supported by ARC DE180100344.

## Notes

### Competing Interest Statement

The authors have declared no competing interest.

